# Structural, dynamic, and evolutionary determinants of substrate binding in the tetrameric 6-phosphogluconate dehydrogenase from *Gluconobacter oxydans*

**DOI:** 10.64898/2026.01.21.700892

**Authors:** Pablo Maturana, Pablo Villalobos, Pietro Roversi, Ricardo Cabrera

**Author notes:** Present address: Plant Biology Department, University of California, Davis, USA. Correspondence to: Pablo Maturana; Ricardo Cabrera.

## Abstract

6-Phosphogluconate dehydrogenases (6PGDHs) catalyze a key oxidative step in the oxidative pentose phosphate pathway (oxPPP), a route essential for NAD(P)H generation and carbon metabolism in bacteria and eukaryotes. While the structural basis of substrate recognition is well established for long-chain dimeric 6PGDHs, the mechanisms used by short-chain tetrameric enzymes remain poorly defined. Here, we present a 2.0 Å crystal structure of tetrameric 6PGDH from *Gluconobacter oxydans* (*Go*6PGDH) in complex with 6-phosphogluconate (6PG) and integrate it with evolutionary, computational, and functional analyses. The structure shows that, unlike dimeric homologs, tetrameric *Go*6PGDH does not undergo a domain-closure transition upon ligand binding. Instead, 6PG induces a compaction of the tetramer mediated by two conserved C-terminal elements: an inter-protomer ionic “lock” and an intra-subunit C-terminal “latch” that together stabilize a closed catalytic pocket. Molecular-dynamics simulations identify His328 as a central residue that couples C-terminal tail closure to direct ligand coordination, and mutagenesis analysis confirms its essential role in catalytic efficiency. Thermodynamic measurements reveal that 6PG binding is strongly enthalpy-driven, consistent with the formation of an ordered hydrogen-bonding and electrostatic network in the closed conformation. These findings define a substrate-induced quaternary-tightening mechanism unique to tetrameric 6PGDHs and illustrate how a conserved C-terminal module has been adapted across the family to regulate substrate binding and catalysis.

## 1. Introduction

The oxidative pentose phosphate pathway (oxPPP) is a central metabolic route crucial for cellular metabolism, supplying reducing power in the form of NAD(P)H and providing ribose-5-phosphate for nucleotide biosynthesis [1]. Within oxPPP, 6-phosphogluconate dehydrogenases (6PGDHs) catalyze the oxidative decarboxylation of 6-phosphogluconate (6PG) to ribulose 5-phosphate (R5P), using NAD⁺ or NADP⁺ as cofactors [2]. Members of the 6PGDH family exist in two oligomeric states, dimeric and tetrameric, which differ in subunit architecture and evolutionary distribution [2,3]. Dimeric enzymes, commonly found in eukaryotes and bacteria, are composed of long-chain subunits containing N- and C-terminal domains. The N-terminal domain adopts a Rossmann fold that binds the cofactor. The C-terminal domain, instead, contains a duplicated set of six alpha helices (hereafter referred to as hemi-domain) that form the 6PG binding site [3]. In contrast, tetrameric enzymes, predominantly found in the bacterial and archaeal lineages [3], consist of short-chain subunits with N- and C-terminal domains, whose C-terminal domains comprise only a single hemi-domain. This structural simplification has led to the proposal that tetrameric 6PGDHs represent evolutionary intermediates between tetrameric β-hydroxy acid dehydrogenases (β-HADH) and the more complex dimeric 6PGDHs [3]. According to this model, structural rearrangements in β-HADHs allowed the C-terminal α-helix to extend across adjacent subunits to form the 6PG binding site in tetrameric 6PGDHs. Subsequent duplication of this hemi-domain gave rise to the dimeric fold, preserving substrate-binding interactions [3].

Structural and mechanistic insights into substrate recognition are well established for dimeric 6PGDHs. Multiple crystal structures of dimeric enzymes in complex with 6PG have shown that the C-terminal region plays a critical role in substrate binding [4–7]. Positively charged residues on the C-terminal α-helix and tail interact with the phosphate and hydroxyl groups of 6PG, stabilizing the substrate through a network of hydrogen bonds and ionic interactions [4,5,7]. For instance, structures of sheep and *Escherichia coli* 6PGDH bound to 6PG reveal that the inter-subunit C-terminal α-helix is a key element in the conformation of of the active site [4,5]. Consistently, studies in *Saccharomyces cerevisiae* demonstrated that deletions in this region abolish enzymatic activity without disrupting the dimeric state, underscoring its catalytic function [8]. Further research into dimeric 6PGDHs has elucidated specific conformational changes triggered by substrate binding. Chen et al. (2010) showed that NADPH binding to one subunit induces a 10° rotation and a 7 Å shift in the coenzyme-binding domain, leading to a closed catalytic conformation [5]. Similarly, Haeussler et al. (2018) reported that in *Plasmodium falciparum* 6PGDH, a flexible loop near the active site rearranges upon 6PG binding, thereby regulating the positioning of the NADP⁺ [6]. In the absence of 6PG, this loop adopts an open conformation, placing NADP⁺ in a “waiting” state. Together, these findings indicate that substrate binding is tightly coupled to conformational changes required for catalysis in dimeric 6PGDHs.

In contrast, for tetrameric 6PGDHs, the structural basis of 6PG recognition remains poorly understood. Studies on *Mycobacterium tuberculosis* 6PGDH *have* identified positively charged residues in the C-terminal α-helix and tail as relevant to catalytic activity [9], yet how this region contributes to substrate binding within the tetrameric assembly remains unclear. To date, no crystallographic structure of a short-chain 6PGDH bound to 6PG has been reported. As a result, it is unknown how tetrameric enzymes organize their substrate-binding pocket, whether ligand binding induces quaternary or intramolecular rearrangements, or how the C-terminal region contributes to catalytic-site assembly in tetrameric 6PGDHs. This lack of structural information hampers our understanding of the evolutionary divergence between dimeric and tetrameric 6PGDH and limits our generalization to generalize substrate-recognition principles across the family.

Here, we address these questions by integrating x-ray crystallography, phylogenetic analysis, molecular dynamics simulations, and functional assays to elucidate the substrate-recognition mechanism of tetrameric 6PGDH from *Gluconobacter oxydans*. The 2.0 Å structure of the enzyme in complex with 6PG reveals that tetrameric 6PGDHs do not undergo a classical domain-closure mechanism upon ligand binding. Instead, substrate binding promotes a C-terminal–mediated quaternary tightening by two structural features: an inter-protomer “lock” and an intra-subunit “latch” that together stabilize a closed catalytic conformation. Evolutionary analysis reveals that the C-terminal α-helix and tail constitute a conserved module in both dimeric and tetrameric 6PGDHs, which becomes progressively extended and more complex in eukaryotic lineages, implying regulation of enzymatic function. Molecular dynamics simulations and mutagenesis further pinpoint His328, a strictly conserved residue in 6PGDH family, as a key residue that couples C-terminal tail closure to catalytic function.

## 2. Materials and Methods

### 2.1. Protein Expression and Purification

Recombinant expression and purification of both wild-type and mutant *Go*6PGDH (Uniprot Entry G5EBD7, 6PGDH_GLUOX) were performed as described by Maturana et. al. [3]. *E. coli* strain BL21(DE3) carrying the expression plasmid of *Go*6PGDH-wild type or *Go*6PGDH^H328A^ was inoculated in Luria-Bertani culture medium at 37 °C. When the cell cultures reached 0.4-0.5 AU at 600 nm, overexpression was induced by adding 1 mM isopropyl-β-D-thiogalactoside (US Biological Swampscott, MA, USA). After 16 hours at 28 °C, the biomass was collected by centrifugation (10 minutes, 4500 × g, 4 °C) using a Thermo Scientific Sorvall RC 6 Plus Centrifuge. Cell extracts were loaded onto a 5 ml HisTrap HP column (GE Healthcare, Chalfont, UK) and equilibrated with the resuspension buffer:50 mM Tris-HCl, 500 mM NaCl, 40 mM imidazole, pH 8.0. For elution, we used a continuous flow with a linear imidazole gradient from 40 mM to 500 mM. The enzymes were purified by size-exclusion chromatography as the second step, using a 16/60 Sephacryl S-200 column (GE Healthcare, Chalfont, UK) in a buffer containing 50 mM Tris-HCl and 100 mM NaCl, pH 8.0. The eluted proteins were concentrated and stored at −20°C.

### 2.2 X-ray crystallography

*Go*6PGDH was concentrated to 12 mg/ml in Tris-HCl buffer (25 mM, pH 8.0), 100 mM NaCl, and 10 mM 6PG. Sitting drop vapor diffusion experiments were performed by mixing 0.8 µL of protein solution and 0.8 µL of crystallization condition, and equilibrating against over 70 µL of mother solution at 18 °C. After optimization, the crystallization condition contained 0.1 M BIS-TRIS at pH 6.5 and 28% w/v polyethylene glycol monomethyl ether 2,000. For cryoprotection, the same solution was supplemented with 20% (v/v) glycerol, and the sample was cooled by plunging it into liquid nitrogen. Diffraction data were collected at beamline 23-ID-B of the Advanced Photon Source, APS (Chicago, IL, USA). Data were indexed and integrated with the XDS software [10], scaled, and merged with Aimless [11], included in the CCP4 suite version 7.1 [12]. Phasing was performed by molecular replacement with MOLREP [13] using the apo structure of *Go*6PGDH (PDB entry 6XQE). The model was iteratively refined using Coot 0.9.8.96 [14] and Phenix Refine from Phenix 1.21.2 [15]. Phenix Refinement was carried out using combined reciprocal-space and real-space coordinate refinement. Local non-crystallographic symmetry (NCS) restraints were applied using a torsion-angle configuration. Atomic displacement parameters were refined as individual isotropic B-factors, and TLS parameters were refined using automatically determined TLS groups. Secondary-structure restraints and automatic optimization of X-ray/stereochemistry weights were applied. Model geometry and stereochemistry were validated using MolProbity software [16].

### 2.3 Structural and sequence analysis

Tetrameric short-chain and dimeric large-chain 6PGDH from *Gluconobacter oxydans* and *Escherichia coli,* respectively, were used as queries to retrieve homologs sequences using BLASP in the clusteredNR database from NCBI. Sequences were aligned using ClustalW with the Blosum62 matrix [17]. A phylogenetic tree of the 6PGDH family was constructed using the maximum-likelihood method implemented in the PhyML server [18]. The VT evolutionary model [19] was selected based on the Akaike Information Criterion (AIC) with the Smart Model Selection (SMS) algorithm [20]. Structural visualization was performed using UCSF Chimera [20] and VMD (Molecular Visual Dynamics, University of Illinois, USA) [21].

### 2.4 Molecular dynamics

Molecular dynamics (MD) simulations were conducted on the tetrameric forms of *Go*6PDH apo (PDBid: 6XQE) and holo (PDBid: 10GW), in the absence of the cofactor. Missing residues in the apo structure were modelled in Modeller 10.6 [22] using the holo structure as a template. The histidine residues in the active site were protonated according to the pKa values determined in the Propka 2.0 server [23]. Proteins were solvated in TIP3P water boxes, and the charges were neutralized by adding Cl⁻ counter ions using AmberTools 24 [24]. Molecular dynamics simulations were performed with AMBER 24 [25], employing the FF19SB protein force field [26] and GAFF2 for 6-phosphogluconate (6PG) available in Amberools 24 [24]. The particle-mesh Ewald method was used to compute electrostatic interactions, with a nonbonded cutoff of 10 Å. The minimization protocol involved 1000 steepest-descent steps followed by 500 conjugate-gradient steps. Three independent simulations were conducted with a timestep of 2.0 fs, during which the systems were gradually heated to 310 K and equilibrated under constant number of particles, pressure, and temperature (NPT ensemble) at 1 atm for 50 ns to ensure stabilization of the tetrameric assembly prior to the production phase. Production simulations were run in triplicate for 200 ns, for a total of 600 ns of cumulative sampling, with a frame saved every 10 ps. RMSD and RMSF Trajectory analysis was performed using the Cpptrajj program from AmberTools 24 [27]. Interactions with 6PG were quantified using an in-house script that uses Prolif 2.9.0 and MdAnalysis 2.9.0. Briefly, for each frame, interactions with 6PG were quantified for each subunit of the tetramer over the three replicates for each system. To quantify residue-residue distances, the overall trajectories were analyzed in VMD by measuring distance using a single atom as a reference for each residue. After that, distance distributions were analyzed using 50 bins per simulation, spanning the minimum to the maximum distance between the two residues. The K-means algorithm in the Cpptraj program was used to identify five protein backbone clusters in the trajectories.

### 2.5 Site-directed mutagenesis

The H328A mutant of *Go*6PGDH was generated using the QuikChange® Site-Directed Mutagenesis Kit (Stratagene) with the plasmid pET-TEV-28a containing *Go*6PGDH cDNA as the template. After PCR amplification, the reaction product was digested with 1U of DpnI (ThermoFisher), transformed into DH5 chemocompetent cells, and cultured in LB-agar medium at 37 °C overnight. Three clones were inoculated in LB at 37°C overnight, and the plasmid DNA was purified using an E. Z. N. A. plasmid DNA mini kit. Plasmids were sequenced by Macrogen (MD, 20850, United States).

### 2.6 Enzyme Kinetics experiments

We measured enzyme activity at room temperature using a BioTek Synergy II spectrophotometer by monitoring absorbance changes at 340 nm associated with NADH formation. Initial rates were obtained from the linear regions of the reaction progress curves. Substrate saturation curves for 6PG were recorded for the wild-type and H328A mutant enzymes in 25 mM Tris–HCl (pH 7.8) and 100 mM NaCl, in a final volume of 0.03 mL. Kinetic parameters were determined by varying 6PG concentration at a fixed NAD^+^ concentration (5 mM). The Michaelis-Menten model: 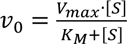, was fitted to the initial velocity data by nonlinear regression using GraphPad Prism 5.0. Best-fit estimates of K_M_ and V_max_ were obtained from the model fitting, and kinetic constants *k_cat_* and k_cat_/K_M_ were derived using the total enzyme concentration in the reaction.

### 2.7 ITC measurements

Before each experiment, 20 mg of total *Go*6PGDH was loaded onto a size-exclusion chromatography column using a preparative column (16/60 Sephacryl S-200 GE) in 50 mM HEPES buffer (pH 7.5) and TCEP (0.5 mM). Titrated and concentrated 6PG (Millipore Sigma Aldrich) was diluted in size-exclusion chromatography buffer. For the 6PG titration, 80 µM *Go*6PGDH was placed in a stirred cell and titrated with 32 injections of 6 µL of 6PG at 1.5 mM each, at 300 s intervals. All measurements were performed at 25 °C by using a VPT-ITC microcalorimeter (Microcal, Northampton, MA, USA). All measurements were performed at 25 °C using a VP-ITC microcalorimeter (MicroCal, Northampton, MA, USA). The raw thermograms were integrated and baseline-corrected in Origin™ (MicroCal), and a single-site binding model was fit to the integrated heat data by nonlinear regression to determine the binding stoichiometry (N) and dissociation constant (K_d_). Thermodynamic parameters (ΔH and ΔH) were derived from the fitted model.

## 3. Results

### 3.1 Structural Insights into Tetrameric Assembly and Active Site Formation of Go6PGDH

To establish the structural basis for substrate recognition in tetrameric 6PGDHs, we determined the crystal structure of *Go*6PGDH in complex with 6PG at 2.0 Å resolution (PDBid: 10GW and Supplementary Table S1). The asymmetric unit includes a homotetramer, with each subunit composed of two domains: an N-terminal Rossmann-fold domain and an all-α-helix C-terminal domain (Fig. 1A). The final α-helix (α14) extends outward from the core of the C-terminal domain, contributing to the active site of an adjacent subunit. The tetramer assembles as two antiparallel dimers stabilized by interactions between the α9 helices (Fig. 1B). Notably, the primary dimer interface in tetrameric 6PGDHs resembles that of the entire C-terminal domain in dimeric 6PGDHs [3,9]. The α14 helix adopts an outward orientation, a feature conserved among tetrameric 6PGDHs, as previously described for *Gluconacetobacter diazotrophicus* 6PGDH [28]. The tetramer is further stabilized by interactions between α14 helices and the hinge regions connecting the N- and C-terminal domains of neighboring subunits. This architecture positions two active sites on opposite faces of the assembly (Fig. 1C).

**Figure 1.**
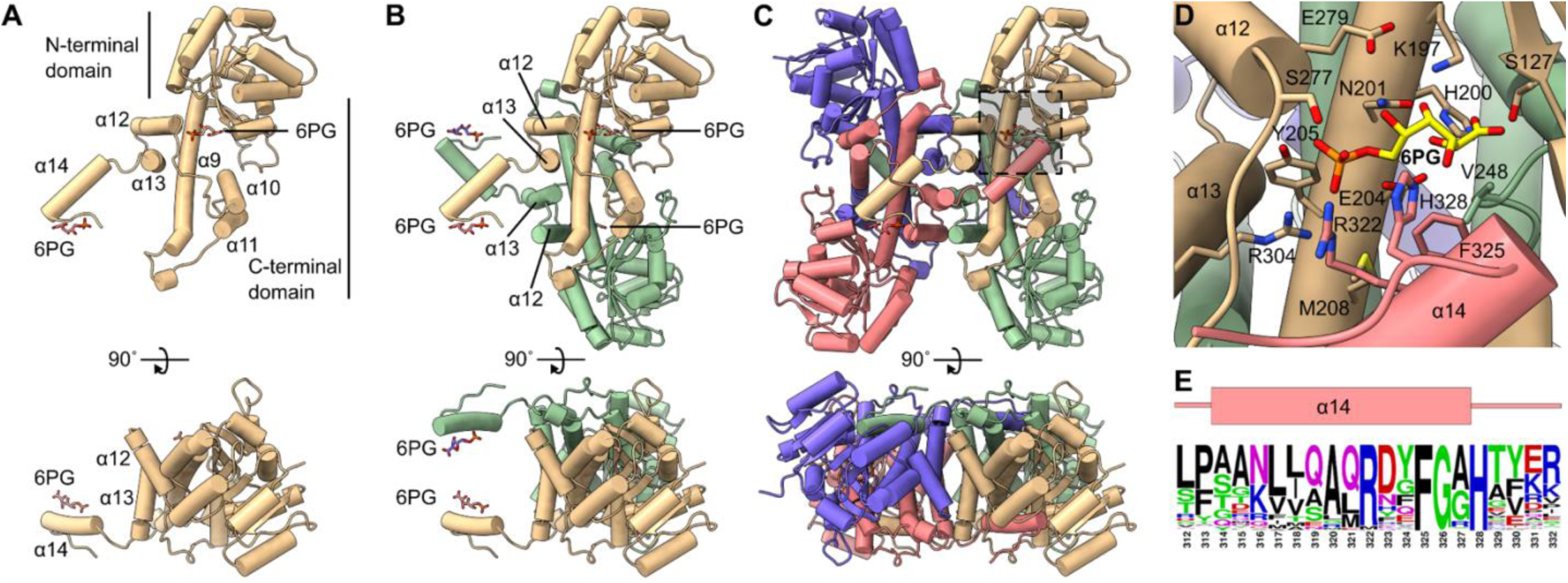
Structural organization and active-site architecture of *Go*6PGDH in complex with 6PG. (A) Monomeric structure of *Go*6PGDH highlighting the N-terminal Rossmann-fold domain and the all-α-helical C-terminal domain. The terminal α-helix (α14) protrudes from the C-terminal domain and extends toward the neighboring subunit, contributing to the formation of the active site. (B) Protomer assembly of *Go*6PGDH illustrating the antiparallel interactions between α9 helices and the relative positioning of α12, α13, and the outward-oriented α14 helices. (C) Tetrameric assembly of *Go*6PGDH composed of two antiparallel dimers (protomers). Subunits are colored individually to emphasize quaternary architecture. Two active sites are positioned on each face of the tetramer, generated at the interfaces between neighboring subunits. (D) Close-up view of the catalytic pocket showing 6PG bound within a groove formed by residues from the N-terminal domain and the α9 and α14 helices of adjacent subunits. Key residues involved in substrate binding (S127, K197, N201, E204, Y205, R304, R322, H328, and F325) are shown as sticks. (E) Sequence conservation of the α14 helix across dimeric and tetrameric 6PGDHs. The sequence logo highlights the conserved RX₂FGXH motif, including residues R322 and H328.

The structure of the complex with 6PG shows the substrate located in a groove formed by the α9 helix of one subunit, residues from the N-terminal domain, and the α14 helix of an adjacent subunit (Fig. 1D and Supplementary Fig. S1). Key residues involved in substrate binding include Ser127, which interacts with the first hydroxyl group of 6PG, and Lys197, Asn201, Glu204, and Tyr205, which create an extended hydrogen-bonding and electrostatic network that anchors the substrate. The β-hydroxyl group of 6PG is positioned 2.9 Å from the amine group of catalytic Lys197, enabling effective catalysis [2,29]. Similar to dimeric 6PGDHs, the interaction with 6PG involves contributions from the terminal α14 helix of the adjacent subunit. Among these residues, Arg322 and His328, located on the α14 helix and the adjacent C-terminal tail, respectively, make direct contacts with the phosphate and γ-hydroxyl groups of 6PG, highlighting the functional relevance of the C-terminal end in shaping the catalytic site. Sequence analysis reveals that the key residues that contact 6PG are highly conserved across members of the 6PGDH family (Fig. 2). Specifically, the α14 helix and the adjacent C-terminal tail harbor the conserved residues Arg322 and His328, which anchor the phosphate and γ-hydroxyl groups of the substrate. The sequence logo emphasizes a conserved C-terminal motif (RX₂FGXH) present in both tetrameric and dimeric enzymes, indicating that this region has remained an essential structural element for 6PG recognition throughout the 6PGDH family (Fig.1E). These observations suggest that the C-terminal region is not merely structural but acts as a ligand-responsive module whose evolutionary history may provide insight into the emergence of substrate-recognition mechanisms.

**Figure 2.**
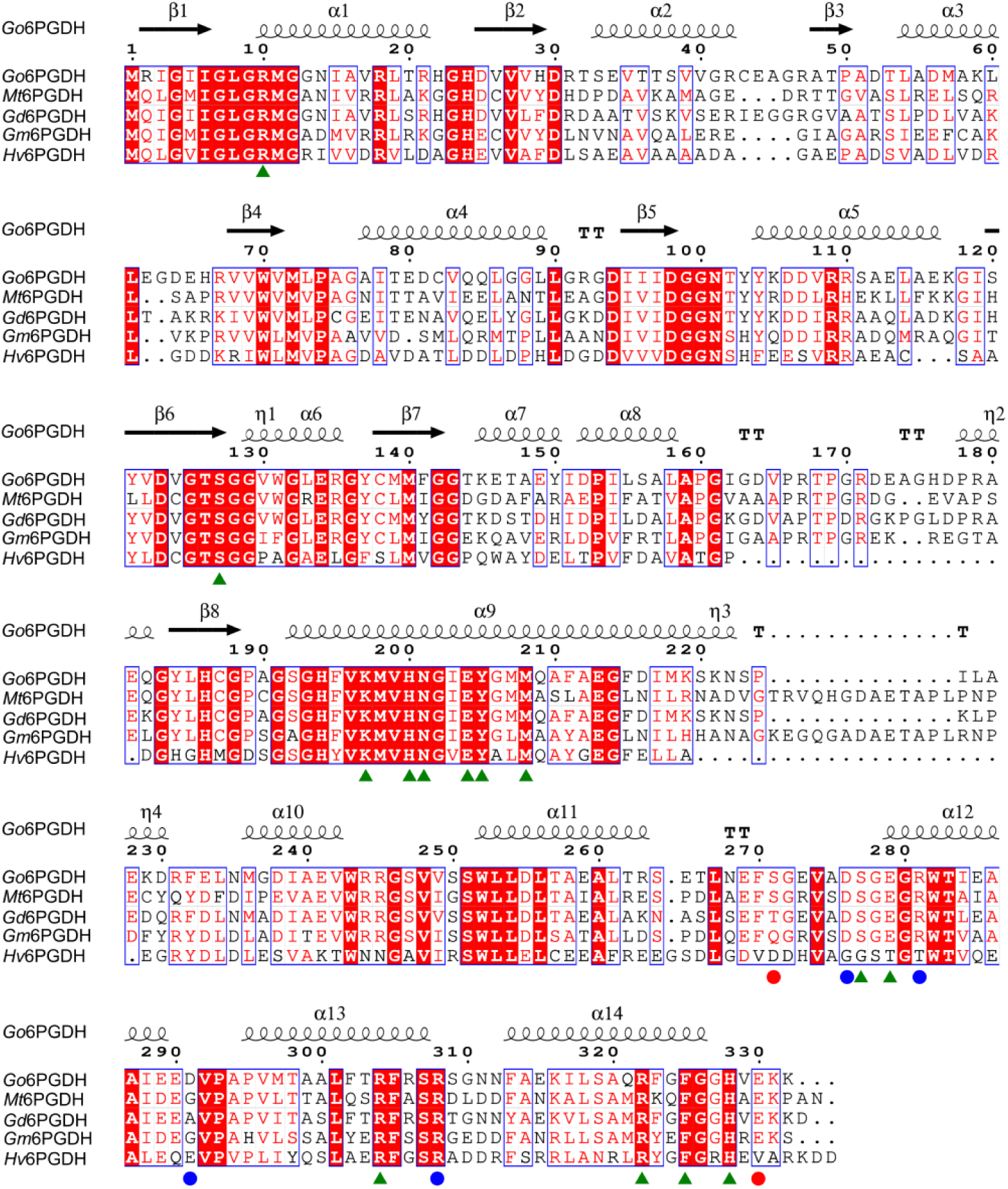
Sequence alignment and conservation of functional residues in tetrameric 6-phosphogluconate dehydrogenases. Multiple sequence alignment of *Gluconobacter oxydans* 6PGDH (*Go*6PGDH) with representative tetrameric homologs, including *Mycobacterium tuberculosis* (*Mt*6PGDH), *Gluconacetobacter diazotrophicus* (*Gd*6PGDH), *Geobacter metallireducens* (*Gm*6PGDH), and *Haloferax volcanii* (*Hv*6PGDH). Secondary-structure elements of *Go*6PGDH are indicated above the alignment. Residues involved in 6-phosphogluconate binding are indicated with green triangles. Residues participating in the inter-protomer “lock” are indicated by blue circles, and residues forming the intra-subunit C-terminal “latch” are highlighted with red circles.

### 3.2 Evolution of the C-terminal end in the 6PGDH Family

Given the important structural role of the C-terminal region in 6PG binding, as described above, we investigated how this region evolved across the 6PGDH family and whether its diversification provides further insight into the factors influencing 6PG recognition. Understanding how the C-terminal region has diversified is crucial for understanding how 6PGDH can modulate substrate binding across different oligomeric forms. To this end, we conducted a phylogenetic analysis to track the evolutionary history of the C-terminal region (Fig. 3A). We also assessed the sequence conservation of this region and performed structural comparisons across representative 6PGDH enzymes. The phylogenetic analysis revealed a clear division between tetrameric and dimeric 6PGDH (Fig. 3A). We found tetrameric enzymes in bacterial and archaeal lineages, forming distinct groups such as Firmicutes, Chloroflexi, and α-Proteobacteria. In contrast, dimeric 6PGDHs are found within bacterial, archaeal, and eukaryotic lineages, including Plants, Euglenozoa, Opisthokonta, and Ascomycetes. The widespread presence of these enzymes across various bacterial and archaeal groups suggests multiple lateral gene transfer events, as previously described by our group [3].

**Figure 3.**
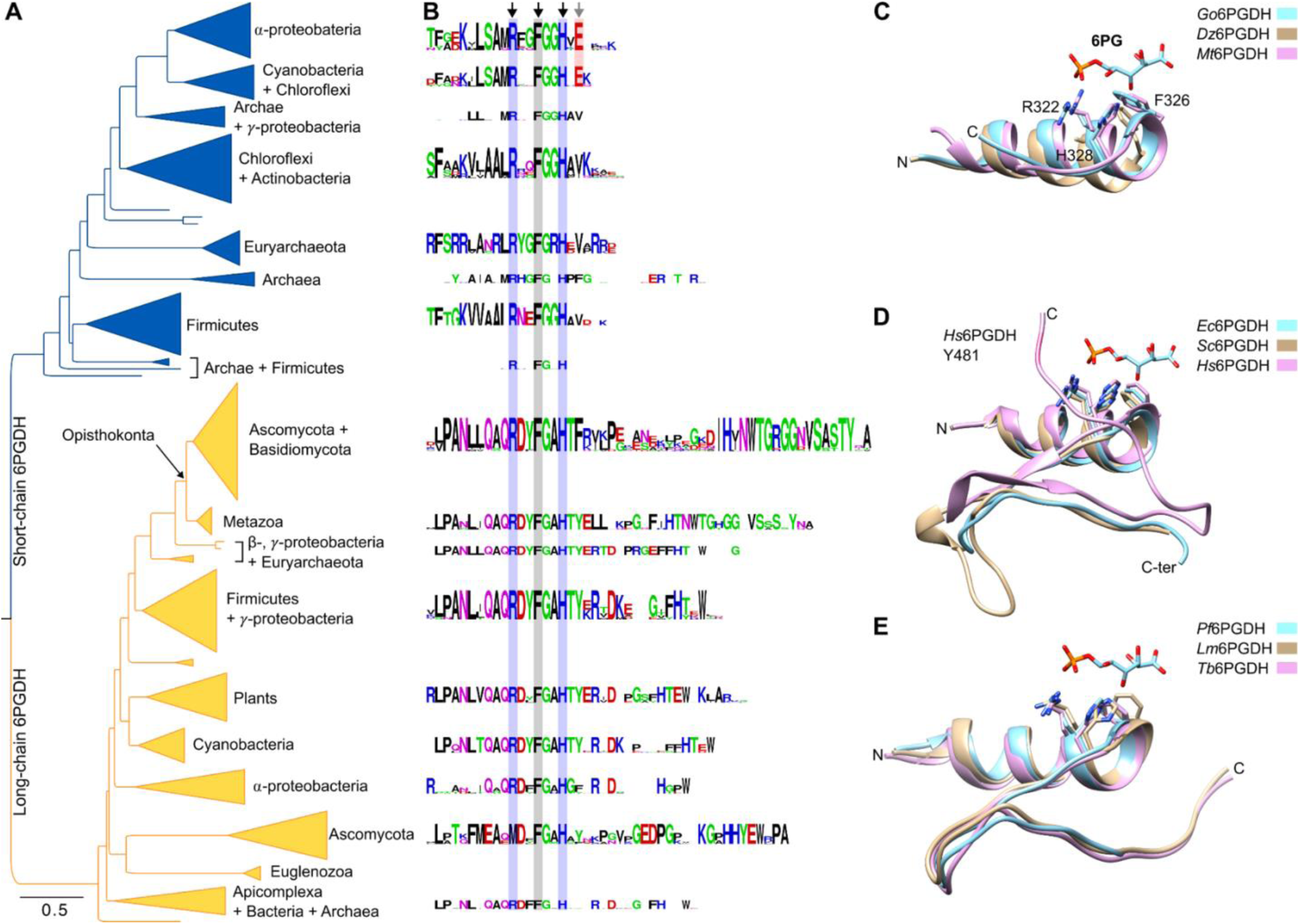
Evolutionary diversification of the C-terminal region in the 6PGDH family. **(A)** Maximum-likelihood phylogenetic tree of the 6PGDH family showing a separation between short-chain tetrameric enzymes (blue) and long-chain dimeric enzymes (yellow). Major taxonomic clades are indicated. This division reflects distinct evolutionary trajectories of the C-terminal region. (B) Sequence logos of the C-terminal region aligned with the phylogenetic clades shown in panel A. Shaded columns highlight conserved positions corresponding to the RX₂FGXH motif. (C) Structural superposition of the C-terminal α14 and tail from representative short-chain 6PGDHs: *Gluconobacter oxydans* (cyan) (PDBid: 10GW), *Gluconoacetobacter diazotrophicus* (beige) (PDBid: 6VPB), and *Mycobacterium tuberculosis* (pink) (PDBid: 9IJB). Conserved residues involved in substrate binding, including R322, F326, and H328 (Go6PGDH numbering), adopt similar spatial positions across bacterial tetramers. (D) Structural comparison of long-chain bacterial and eukaryotic 6PGDHs, including *Escherichia coli* (cyan; 6PG-bound) (PDBid: 3FWN), *Saccharomyces cerevisiae* (beige) (PDBid: 2P4Q), and human 6PGDH (pink) (PDBid: 4GWG). These enzymes show progressively extended C-terminal regions that project toward the catalytic pocket. In human 6PGDH, Y481 is oriented toward the substrate-binding site. (E) Structural superposition of euglenozoan 6PGDHs (*Plasmodium falciparum*, cyan (PDBid: 6FQZ); *Leishmania donovani*, beige (PDBid: 8C79); *Trypanosoma brucei*, pink (PDBid: 1PGJ)), illustrating an intermediate C-terminal architecture between bacterial short-chain enzymes and Opisthokont long-chain homologs.

To explore the structural implications of the C-terminal end, we analyzed its sequence motif across various clades (Fig. 3B). In short-chain 6PGDHs, the RX₂FGXH motif within the α14 helix is highly conserved and essential for substrate binding. In *Go*6PGDH, residues R322 and H328 stabilize the phosphate and hydroxyl groups of 6PG, a feature conserved in other short-chain enzymes such as *Mycobacterium tuberculosis* [9] and *Gluconoacetobacter diazotrophicus* [28] (Fig. 3C). These enzymes possess a short C-terminal region with minor tails extending beyond the α14 helix, which do not interact with the substrate. In long-chain 6PGDHs, the C-terminal tail region shows greater variability and complexity. Superimposed structures of *E. coli* 6PGDH [5], *Saccharomyces cerevisiae* 6PGDH [8], and human 6PGDH [30], highlight progressive elongation of the C-terminal region (Fig. 3D). In Opisthokonta, a second conserved motif (HTNWTGXGGXVSXYXA) appears, creating a tail extension that positions residues like Y481 (in humans) within the cofactor-binding pocket for NADP^+^ [30]. This tyrosine, unique to Opisthokonts, has been involved in post-translational regulation, suggesting a dual role for the extended C-terminal in catalysis and regulation [2,31]. Sequence logo analysis further supports the evolutionary development of this extended motif, which is missing in bacterial and basal eukaryotic groups. Notably, Euglenozoa group, such as *Plasmodium falciparum* [6], *Leishmania donovani* [32], and *Trypanosoma brucei* [33], demonstrate an intermediate state. Their C-terminal regions are longer than those in bacteria but shorter than in Opisthokonts, indicating a transitional evolutionary intermediate (Fig. 3F). These results suggest a gradual extension of the C-terminal domain, with increasing complexity and regulatory functions in advanced eukaryotic lineages.

Overall, our analysis shows an evolutionary path of the 6PGDH C-terminal end, from short motifs in bacterial enzymes to longer regions in Opisthokonts. These extensions could be linked to functional adaptations, such as improved substrate specificity, catalytic efficiency, and post-translational regulation, highlighting the evolutionary diversity of 6PGDH enzymes. The evolutionary diversification of the C-terminal region suggests a tight link between its structure and ligand-dependent responses. We therefore examined how 6PG binding reshapes *Go*6PGDH using comparative structural and MD analyses.

### 3.3 Conformational changes and interactions associated with the 6PG binding in Go6PGDH

To characterize the conformational changes involved in 6PG binding to *Go*6PGDH, we compared its holo (6PG-bound) and apo states and evaluated them in the context of all other tetrameric homologs (*Geobacter metallireducens, Gluconacetobacter diazotrophicus*, and *Mycobacterium tuberculosis*). Structural superposition of all four subunits in holo *Go*6PGDH revealed that they nearly adopt identical fold, with well-aligned N- and C-terminal domains (Fig. S2A). Comparing the holo *Go*6PGDH monomer with the apo structures of *G. metallireducens*, *G. diazotrophicus*, and *M. tuberculosis* 6PGDH showed that the subunit fold is conserved across these tetrameric enzymes (Fig. S2B). However, some structural differences appeared when comparing the holo *Go*6PGDH structure to its apo form. In subunits B and D, the α14 helix showed a 24° outward rotation, pivoting around the α13 helix (Fig. S2C). In contrast, subunits A and C remained aligned with the holo structure. This conformational change is limited to diagonally opposite subunits, leading to a symmetrical arrangement where α14 helices within each pair (A/C and B/D) face the same direction on the tetramer (Fig. S2D). At the quaternary level, the apo assembly exhibits an additional ∼25° inter-dimer rotation, creating a more expanded tetramer compared to the compact holo state (Fig. S2D). Therefore, ligand binding results in a tightening of the quaternary interface rather than significant rearrangements of the N- or C-terminal domains (Fig. S3). This compaction stabilizes the active sites by reducing inter-subunit distances and optimizing the catalytic configuration.

To examine the conformational dynamics related to 6PG binding in *Go*6PGDH, we conducted 200-ns molecular dynamics (MD) simulations in triplicate for both apo and 6PG-bound forms using the AMBER force field. RMSD analysis confirmed that all simulations remained stable (Fig. S4A and B). The trajectories from the production runs were analyzed using RMSF to evaluate differences in flexibility between systems with and without 6PG (Fig. 4A).

**Figure 4.**
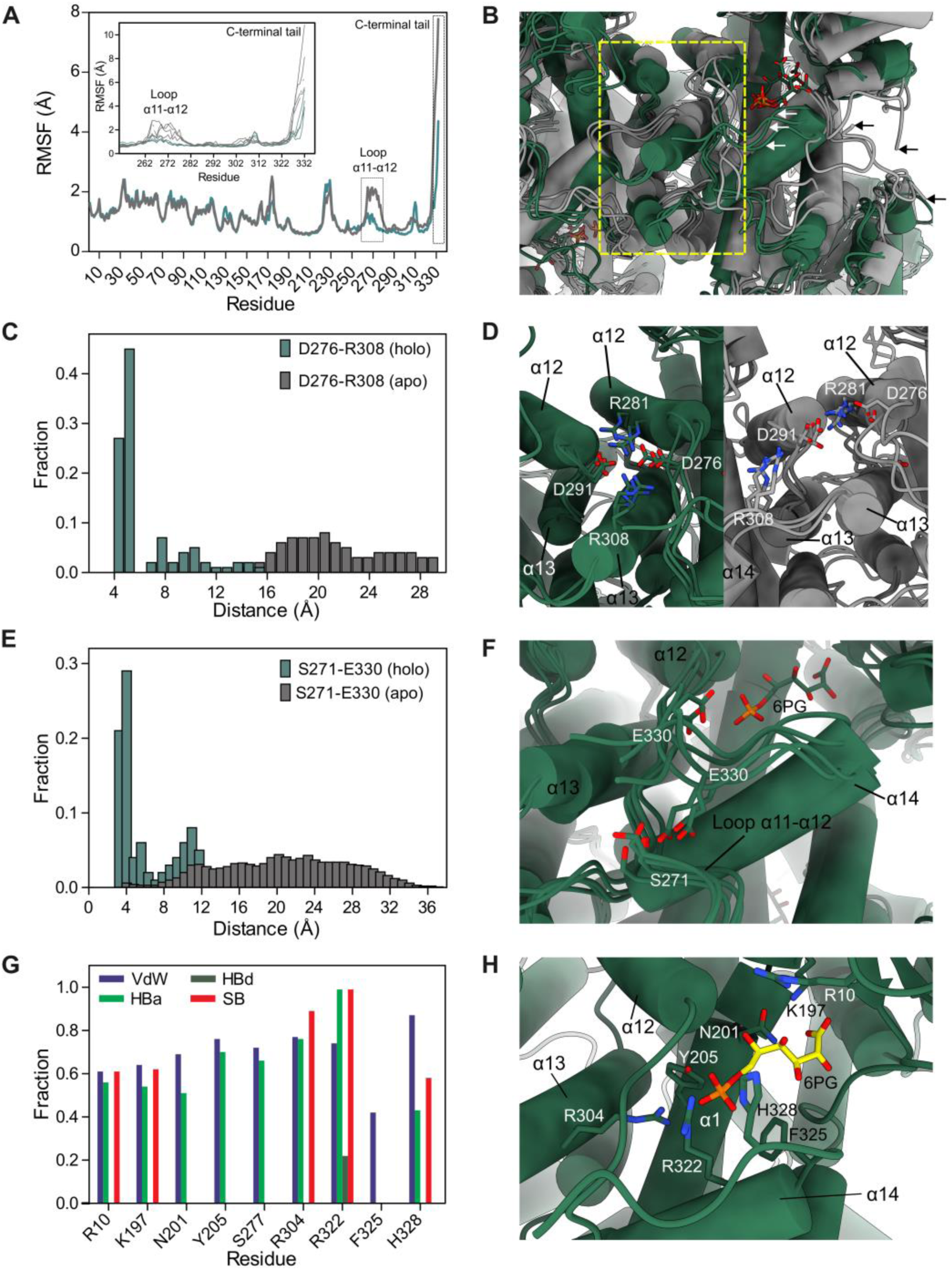
Ligand-induced conformational dynamics and quaternary tightening in *Go*6PGDH. **(A)** Backbone root-mean-square fluctuation (RMSF) profiles obtained from three independent 200-ns molecular dynamics simulations of apo (gray) and 6PG-bound (green) *Go*6PGDH. Substrate binding markedly reduces flexibility in the α11–α12 loop (residues 265-278) and in the C-terminal tail (residues 319-332), as highlighted in the inset. (B) Superposition of representative centroid structures from K-means clustering of the combined apo (gray) and holo (green) trajectories. The dashed box highlights the inter-protomer region undergoing ligand-dependent rearrangement. Apo centroids sample a broader range of conformations for the C-terminal tail (black arrows), whereas holo centroids converge toward a more compact and ordered arrangement around the active site (white arrows). (C) Distance distributions for the inter-protomer D276-R308 residue pair across apo (gray) and holo (green) simulations. (D) Structural comparison of the α12-α13 inter-protomer interface in the holo (left, green) and apo (right, gray) states, illustrating stabilization of the D276-R308 interaction and associated residues (D291, R281) upon ligand binding. This interaction underlies ligand-induced quaternary tightening of the tetramer. (E) Distance distributions for the intra-subunit S271-E330 residue pair, defining the intra-subunit C-terminal “latch”. (F) Structural view of the intra-subunit latch in the holo enzyme, showing E330 from the C-terminal tail forming stabilizing interactions with S271 in the α11–α12 loop for the main three centroids at ∼4Å distance, for other two centroids, E330 is oriented away from S371 (∼11 Å distance). (G) Interaction frequencies between 6PG and surrounding residues within 10 Å, classified by interaction type: van der Waals in purple (VdW), hydrogen-bond acceptor in light green (HBa), hydrogen-bond donor in dark green (HBd), and salt bridge in red (SB. (H) Structural representation of the 6PG-binding site in the holo state, highlighting the network of residues, including R10, K197, N201, Y205, R304, R322, F325, and H328.

The most notable RMSF changes occur in the loop α11-α12, covering residues 265-278, and the C-terminal tail spanning residues 327-332 (inset graph, Fig. 4A). Both regions show increased flexibility in the apo enzyme, suggesting that 6PG binding limits their conformational dynamics. To further explore these rearrangements, we performed cluster analysis on each trajectory (five clusters per state; Supplementary Table S2 and Fig. S4C). Superposition of the representative structure of each cluster (centroids) (Fig. 4B) shows that 6PG stabilizes conformations where the C-terminal tail is closer to the substrate and the 265-278 loop (white arrows, Fig. 4B). Conversely, apo centroids display a wider range of conformations, with both regions adopting open and relaxed states (black arrows Fig. S3C, and Supplementary Table S2).

A key finding from the simulations is the formation of stable interprotomer contacts between Asp276 (from the 265-278 loop), Arg281, and Arg308 from one subunit, and Asp291 from an adjacent protomer (blue circles, Fig 2). We used the intra-subunit interaction Asp276-Arg308 to quantify the formation of these interactions (Fig. 4C). In the 6PG-bound trajectories, the D276-R308 distance distribution is centered at 4-6 Å, consistent with stable ionic or hydrogen-bonding interactions, and shows minimal sampling at larger distances (Fig. 4C and D). This distribution reflects that, in most tetrameric interfaces, Asp276 and Arg308 remain engaged as part of a conserved interaction network involving Asp276, Arg281, Arg308, and Asp291 (Fig. S4D-G). A minor population at longer distances likely corresponds to a subset of interfaces in which Arg308 becomes transiently disengaged from this network and adopts an alternative orientation, pointing away from Asp276 (Fig. S4D-G). In contrast, the apo enzyme shows little sampling below ∼8 Å and a broad distribution from ∼10 to 28 Å, indicating a disorganized and exposed interface (Fig. 4C and D). This interaction acts as an inter-protomer “lock” that stabilizes the dimer-dimer interface and is directly associated with the tetrameric compaction seen in the holo crystal structure (Fig. 4D). The formation of these ionic interactions offers a mechanistic explanation for how 6PG binding causes quaternary tightening without requiring large intrasubunit domain movements.

A second, more localized conformational change involves the closure of the C-terminal tail over the active site, mediated by the E330-S271 C-terminal “latch” (red circles, Fig. 2). This motion is captured by the distance between Glu330 and Ser271 (Fig. 4E). In the 6PG-bound state, the distribution of E330-S271 is centered around 4-6 Å, with contact analysis showing a hydrogen-bond frequency of 0.6. In the apo simulations, these distances increase significantly, ranging from about 10 to 30 Å with minimal sampling within hydrogen-bonding range, indicating that the tail is fully open and flexible (black arrows Fig. 4B). Along with His328, which secures the C-terminal tail against the ligand, E330 functions as an intra-subunit latch that stabilizes the closed conformation only when the substrate is present (white arrows Fig. 4B and Fig. 4F). The strict conservation of E330 across α-proteobacterial, cyanobacterial, and Chloroflexi tetrameric 6PGDHs highlights its functional significance as a defining structural element that mediates catalytic-site closure within these groups (Fig. 3B).

Having established the global and local rearrangements associated with 6PG binding, we next quantified residue-ligand interaction frequencies within the catalytic pocket. Proximity and contact-type analyses of residues within 10 Å of 6PG (Fig. 4G-H) further demonstrate how substrate binding reinforces this network. Most residues contact 6PG primarily through van der Waals (VdW) and hydrogen-bond acceptor (Hba) interactions, forming a broad stabilizing network around the ligand. Notably, Arg322 and His328 stand out. Arg322 shows the highest overall interaction frequencies and is the only residue that engages in all four interaction types: van der Waals, hydrogen-bond acceptor, hydrogen-bond donor (HBd), and salt-bridge interactions (SB), indicating a central electrostatic and hydrogen-bonding role in coordinating the phosphate group. His328 exhibits high VdW and HBa occupancies along with a strong salt-bridge component, reflecting its versatile chemical engagement with both the phosphate and γ-hydroxyl groups of 6PG. Other pocket residues (e.g., Arg10, Lys197, Asn201, Tyr205, Ser277, Arg304) primarily contribute through VdW and HBa interactions, with SB contacts established with Arg10 and Lys197, and Arg304 interacting with the phosphate group of 6PG, providing additional stabilizing contacts but without the diverse interaction modes observed for Arg322 and His328. Phe325 interacts solely via VdW contacts, reflecting its hydrophobic position beneath the C-terminal α14 helix. This interaction pattern is consistent with the conservation of key residues at the site: Arg322 is retained across most members of the family (except in Ascomycota dimeric 6PGDHs, Fig. 3B), while His328 instead, is the only C-terminal residue with strict conservation pattern in both dimeric and tetrameric enzymes (Fig. 3B). Positioned at the base of the C-terminal tail, His328 stabilizes the closed catalytic pocket in the holo state and consistently engages 6PG throughout the structural and MD analyses. These observations guided us to examine its functional relevance directly, leading into the kinetic characterization of the H328A variant described below.

### 3.4 Functional analysis of 6PG binding to Go6PGDH

To elucidate the functional role of H328 in 6PG catalysis, we conducted kinetic assays comparing the wild-type *Go*6PGDH and the H328A mutant (Fig. 5A). Saturation curves for 6PG revealed notable differences between two enzymes. The wild-type *Go*6PGDH showed a Michaelis-Menten constant (K_M_) for 6PG of 0.83 ± 0.08 mM and a turnover number (*k_cat_*) of 47.2 ± 3.5 s^-^¹, resulting in a catalytic efficiency (*k_cat_* /K_M_) of 56.9 ± 6.9 s⁻¹ mM⁻¹ (Fig. 5B). In contrast, the H328A mutant displayed a threefold increase in K_M_ (2.5 ± 0.2 mM) and about a sixfold decrease in kcat (7.4 ± 0.4 s⁻¹), leading to a catalytic efficiency of 3.0 ± 0.3 s⁻¹ mM⁻¹ (Fig. 5B). These findings highlight the critical role of H328 in maintaining substrate affinity and catalytic turnover, as its substitution causes a significant drop in enzyme efficiency.

**Figure 5.**
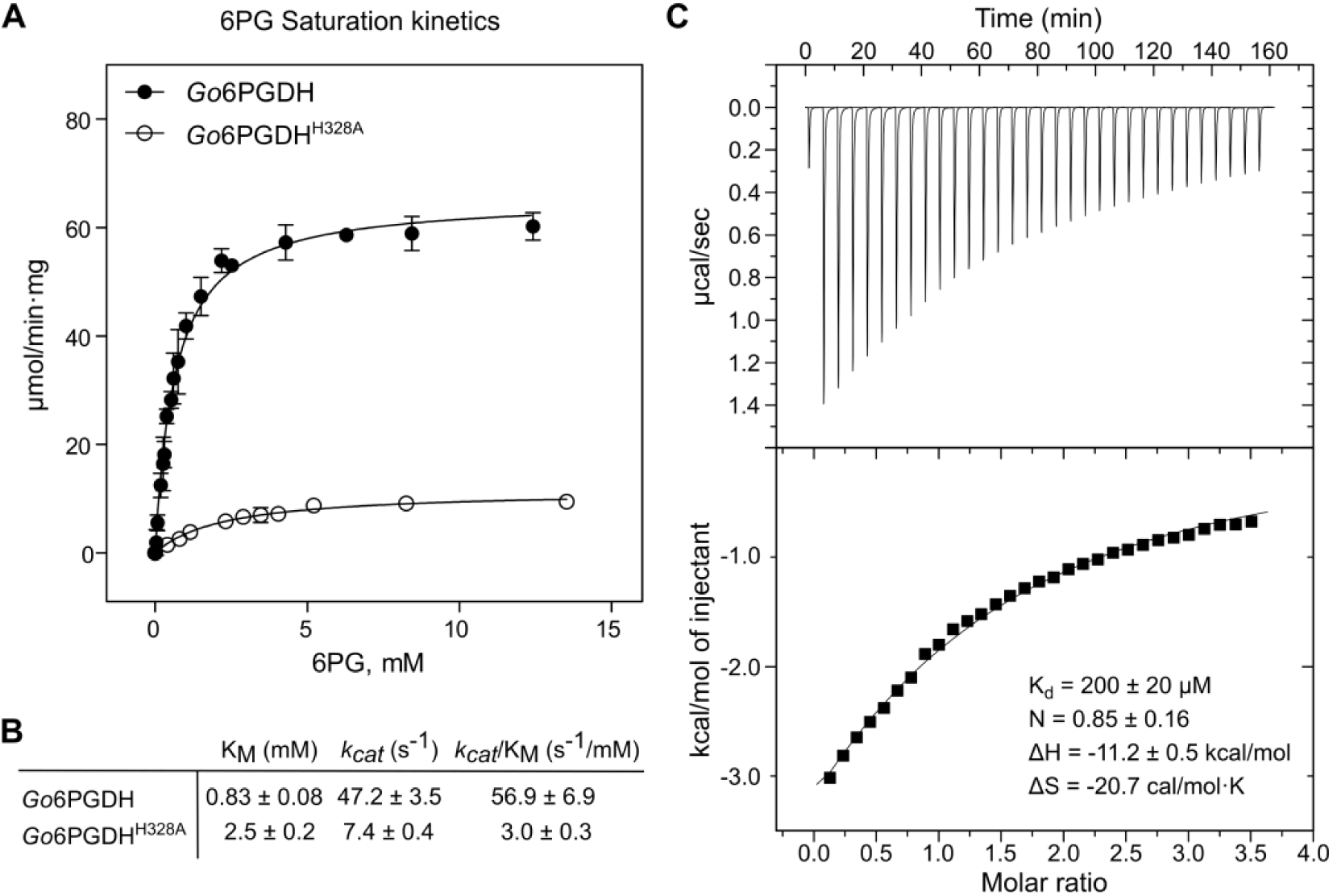
Functional analysis of 6PG binding to *Gluconobacter oxydans* 6PGDH. (A) Kinetic behavior of the wild-type *Go*6PGDH (black circles) and the H328A mutant (white circles). The enzymatic activity was measured as a function of 6PG concentration, with NAD⁺ at 5 mM. Error bars represent the standard deviation from three independent experiments. (B) Kinetic parameters for 6PG of wild-type *Go*6PGDH and *Go*6PGDH^H328A^ enzymes obtained from the kinetic curves. (C) Isothermal titration calorimetry (ITC) analysis of 6PG binding to *Go*6PGDH. The upper panel shows the raw calorimetric data, while the lower panel depicts the integrated binding isotherm. Thermodynamic parameters, including dissociation constant (K_d_), enthalpy (ΔH), and Gibbs free energy (ΔG), indicate a spontaneous, enthalpy-driven binding process with moderate affinity.

Structural and MD data showed that 6PG promotes closure of the catalytic pocket and quaternary tightening, suggesting an underlying energetic profile associated with 6PG binding. To investigate this, we measured 6PG binding using isothermal titration calorimetry (ITC). The ITC results supported the kinetic data by providing thermodynamic details of 6PG binding (Fig. 5C). The binding stoichiometry (N) was found to be 0.85 ± 0.16, suggesting that each subunit of the tetramer generally interacts with one 6PG molecule. The dissociation constant (K_d_) was determined to be 200 ± 20 μM, reflecting a moderate affinity between 6PG and *Go*6PGDH. The binding was exothermic, with an enthalpy change (ΔH) of −11.2 ± 0.5 kcal/mol, and involved an unfavorable entropy change (ΔS) of −20.7 cal/mol·K. The Gibbs free energy change (ΔG) at 298 K was calculated as −5.0 kcal/mol, confirming that the interaction is spontaneous. These thermodynamic parameters indicate that binding is mainly enthalpically driven, likely by hydrogen bonds and ionic interactions at the active site.

Together, the structural, kinetic, and thermodynamic data point to a unified mechanism where His328 is a key factor in recognizing 6PG and enabling catalytic activity. The significant reduction in efficiency observed in the H328A mutant highlights its dual role in stabilizing the closed-pocket conformation and facilitating optimal transition-state formation. These findings align well with the previously described conformational framework, underscoring the importance of interactions between the C-terminal α14 helix and tail in linking substrate binding to catalytic function in *Go*6PGDH.

## 4. Discussion

This study presents the crystal structure of the tetrameric 6PGDH from *Gluconobacter oxydans* in complex with 6-phosphogluconate at 2.0 Å resolution and provides a mechanistic framework that integrates structural, evolutionary, and functional determinants of substrate recognition in short-chain tetrameric enzymes. Our findings reaffirm the conserved fold of the active site across the 6PGDH family while demonstrating that tetrameric enzymes rely on a distinct mode of catalytic-site assembly that diverges fundamentally from the domain-closure mechanism characteristic of long-chain dimeric homologs. Instead of large intramolecular motions, *Go*6PGDH constructs its active site through a composite interface in which the C-terminal α-helix (α14) and the adjacent tail, contributed by the neighboring subunit, close over the substrate. This arrangement positions residues such as Arg322 and His328 in direct proximity to the phosphate and hydroxyl groups of 6PG, underscoring the central role of the C-terminal module in substrate coordination and providing a structural rationale for how short-chain tetramers compensate for the absence of duplicated domains found in long-chain enzymes. Together, these observations suggest that tetrameric 6PGDHs do not represent simplified versions of their dimeric counterparts, but instead implement a distinct strategy for coupling substrate binding to catalytic function.

The crystallographic data confirm that short-chain bacterial 6PGDHs are genuine tetramers and support our earlier proposal, first advanced by our group, that these enzymes represent evolutionary intermediates between β-hydroxyacid dehydrogenases and more derived dimeric 6PGDHs [3]. This structural evidence contradicts an alternative dimeric model, which suggested membrane-anchored dimers stabilized by C-terminal ends interactions. Instead [28], our structure shows a quaternary arrangement in which the C-terminal α-helix is not a membrane anchor but an integral component of the catalytic architecture (Fig. 1). Critical residues within the active site, Ser127, Lys197, Asn201, Glu204, and Tyr205, form an extended hydrogen-bonding network with 6PG, consistent with substrate positioning seen in dimeric enzymes from *E. coli*, *P. falciparum*, and *G. stearothermophilus* [5–7] (Fig. 1C). The conservation of this catalytic scaffold across oligomeric forms highlights evolutionary pressure to preserve core chemistry while allowing diversification of additional regulatory or structural features.

Phylogenetic and structural analyses further reveal that the short C-terminal end carrying the conserved RX₂FGXH motif constitutes a compact, functionally specialized module that has been selectively maintained across α-Proteobacteria, Cyanobacteria, and Chloroflexi. This motif anchors Arg322 and His328 at the substrate-binding interface and appears optimized for the tetrameric fold rather than representing a vestigial remnant (Fig. 1E and 2B). In contrast, Opisthokont dimeric 6PGDHs exhibit an expanded C-terminal end with additional conserved motifs that support regulatory functions likely not required in bacterial tetramers (Fig. 2B). Intermediate C-terminal ends found in Euglenozoa, possessing C-terminal regions of intermediate length, suggest that this diversification occurred stepwise. Thus, what begins as a minimalist substrate-binding scaffold in bacterial tetramers evolves into a versatile regulatory appendage in more complex eukaryotic lineages, indicating that the C-terminal end is a dynamic evolutionary module rather than a static structural tail (Fig. 2B). From an evolutionary perspective, this progressive C-terminal elongation suggests that the ancestral function of the C-terminal module was purely catalytic, with regulatory roles emerging later as metabolic control became more complex in eukaryotic lineages. In this context, tetrameric bacterial 6PGDHs may represent an optimized solution for robust catalysis under fluctuating metabolic conditions, where regulation through additional structural elements is unnecessary or even disadvantageous.

The comparison between holo- and apo-tetrameric states demonstrates that 6PG binding elicits conformational changes that remodel the quaternary structure association without triggering domain closure. Unlike dimeric enzymes, which rely on domain closure to assemble the catalytic pocket, tetrameric *Go*6PGDH undergoes subtler but functionally significant rearrangements. The holo structure exhibits a 24° outward rotation of α14 in two subunits, while inter-dimeric rotation angles contract by ∼25°, generating a more compact assembly. Molecular dynamics simulations explain this compaction through two ligand-stabilized interactions that function cooperatively: a persistent ionic “lock” conformed by Asp276, Arg281, Asp291and Arg308 at the protomer interface (Fig. 4C and 4D), and a C-terminal “latch” involving Glu330 and Ser271 that restricts the mobility of the tail region and positions His328 for optimal engagement with 6PG (Fig. 4E and F). These interactions display narrow distance distributions and prolonged occupancies in the holo state but may remain highly dynamic and solvent-exposed in the apo simulations. This lock-latch mechanism exemplifies an alternative mode of enzyme activation in which ligand binding stabilizes quaternary geometry rather than inducing intramolecular domain rearrangements. Such a strategy may be particularly advantageous for compact oligomeric enzymes, where catalytic regulation must be achieved without the architectural redundancy present in multi-domain enzymes.

Thermodynamic data corroborate this mechanistic interpretation. ITC experiments demonstrate that 6PG binding is enthalpy-driven with a negative ΔH and unfavorable ΔS, indicating the formation of an ordered network of intermolecular contacts upon ligand binding. The conformational tightening mediated by the lock-latch interactions provides a structural basis for these energetic signatures, as increased ordering of inter-protomer interfaces promotes enthalpic stabilization despite entropic cost. This model explains how tetrameric 6PGDHs could achieve high catalytic efficiency through concerted adjustments in quaternary geometry rather than relying on major intramolecular rearrangements. Functional evidence further supports the significance of the C-terminal end. The H328A variant exhibits a marked increase in K_M_ and a substantial reduction in *k_cat_*, resulting in a ∼19-fold loss in catalytic efficiency (Fig. 5B). This result is consistent with the observations of Wang *et al.*, who reported a pronounced reduction in enzymatic activity for the H328A mutant, although a detailed kinetic characterization was not performed [9]. The decrease in *k_cat_* and increase of K_M_ reflect both the loss of direct His328-6PG interactions and an inability to achieve proper catalysis, consistent with the structural and dynamic data (Fig. 4). The mutation, therefore, uncouples substrate binding from productive active-site assembly, underscoring the role of His328 in ligand coordination and in stabilizing the conformational tightening necessary for catalysis.

Our results support a unified model in which tetrameric 6PGDHs employ a conserved C-terminal end to mediate substrate recognition and to orchestrate ligand-induced quaternary tightening. This mechanism reconciles structural simplicity with catalytic efficiency and provides an evolutionary rationale for the strict conservation of the RX₂FGXH motif in bacterial tetramers. More broadly, the lock-latch strategy described here suggests an alternative solution by which compact oligomeric dehydrogenases can achieve substrate-induced activation in the absence of large-scale domain movements. By integrating crystallographic, evolutionary, dynamic, and functional analyses, this study establishes a cohesive framework for understanding substrate recognition and catalytic activation in tetrameric 6PGDH enzymes and lays the groundwork for future investigations into how structural heterogeneity shapes enzyme function across the broader dehydrogenase superfamily.

### CRediT authorship contribution statement

PM: Conceptualization; methodology; investigation; formal analysis; funding acquisition; project administration; writing-original draft; writing-review & editing. PV: Methodology; investigation, formal analysis; software; writing-original draft. PR: Methodology, data curation; software; writing-review & editing. RC: Conceptualization; formal analysis; investigation; supervision; funding acquisition; project administration.

## Data availability

The structure coordinates are available in the Protein Data Bank under accession code 10GW.

## Acknowledgments

We thank Dr. Víctor Castro Fernández for providing the opportunity and support for X-ray diffraction through his research project, and Dr. Felipe González Ordenes for assistance with diffraction data collection. The GM/CA@APSbeamline 23-ID-B staff is acknowledged for their assistance during the diffraction experiments, especially to Ruslan Sanishvili. GM/CA@APS has been funded by the National Cancer Institute (ACB-12002) and the National Institute of General Medical Sciences (AGM-12006, P30GM138396). This research used resources of the Advanced Photon Source, a U.S. Department of Energy (DOE) Office of Science User Facility operated for the DOE Office of Science by Argonne National Laboratory under Contract No. DE-AC02-06CH11357.

## Supplementary material

**Figure S1.**
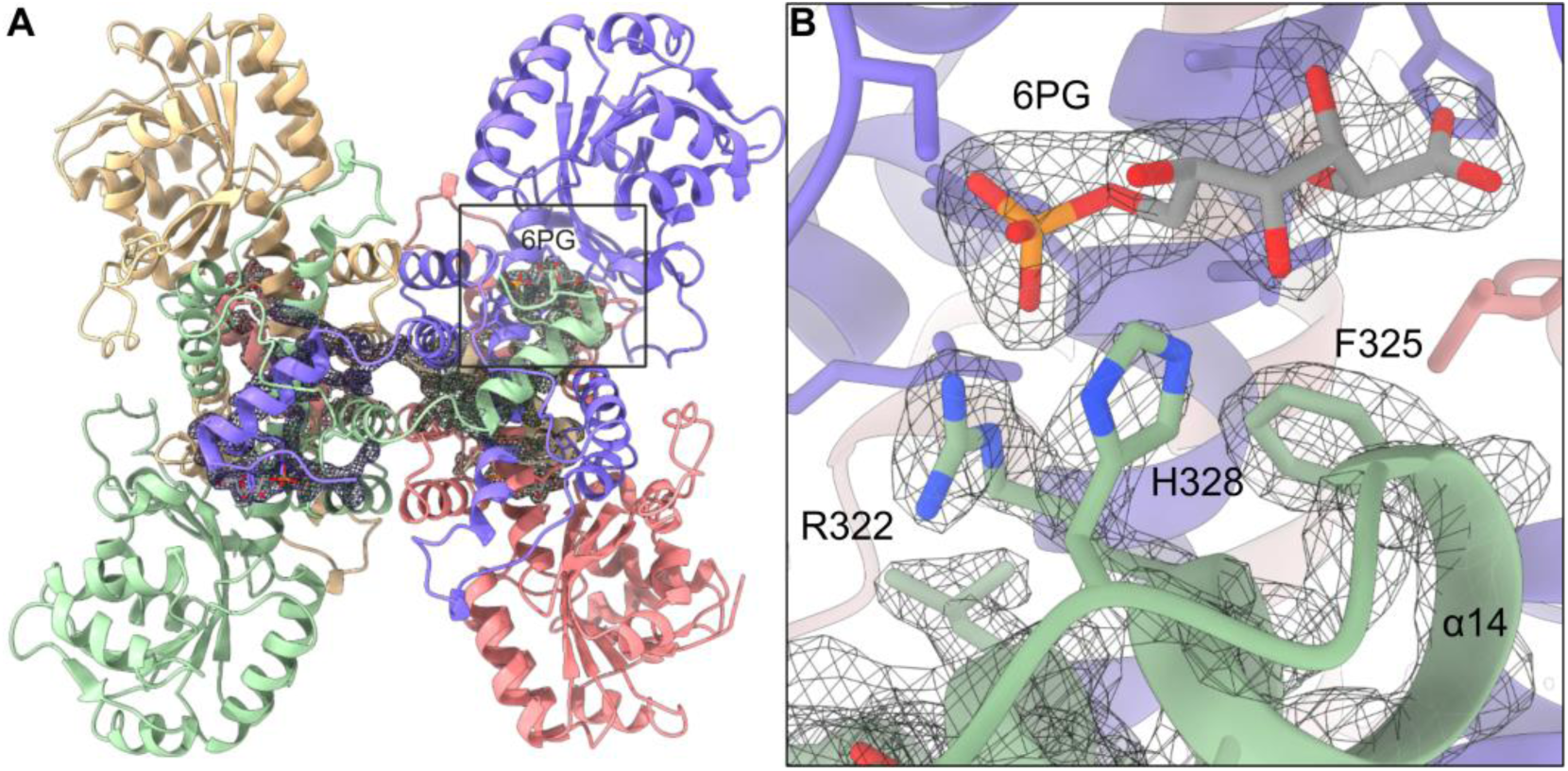
Polder OMIT map for 6-phosphogluconate. (Left) Overall view of the *Go*6PGDH tetramer, highlighting the 6PG-binding pocket. A polder OMIT map of 6PG, and the C-terminal end was calculated to validate ligand placement and is displayed around the active site. (Right) Close-up of the catalytic pocket showing the 6PG ligand and surrounding residues (R322, F325, H328). The polder OMIT map is contoured at 3.0 σ and reveals well-defined positive density for the phosphate and hydroxyl groups of 6PG, confirming unambiguous ligand position and its interactions with the α14 helix and adjacent tail.

**Figure S2.**
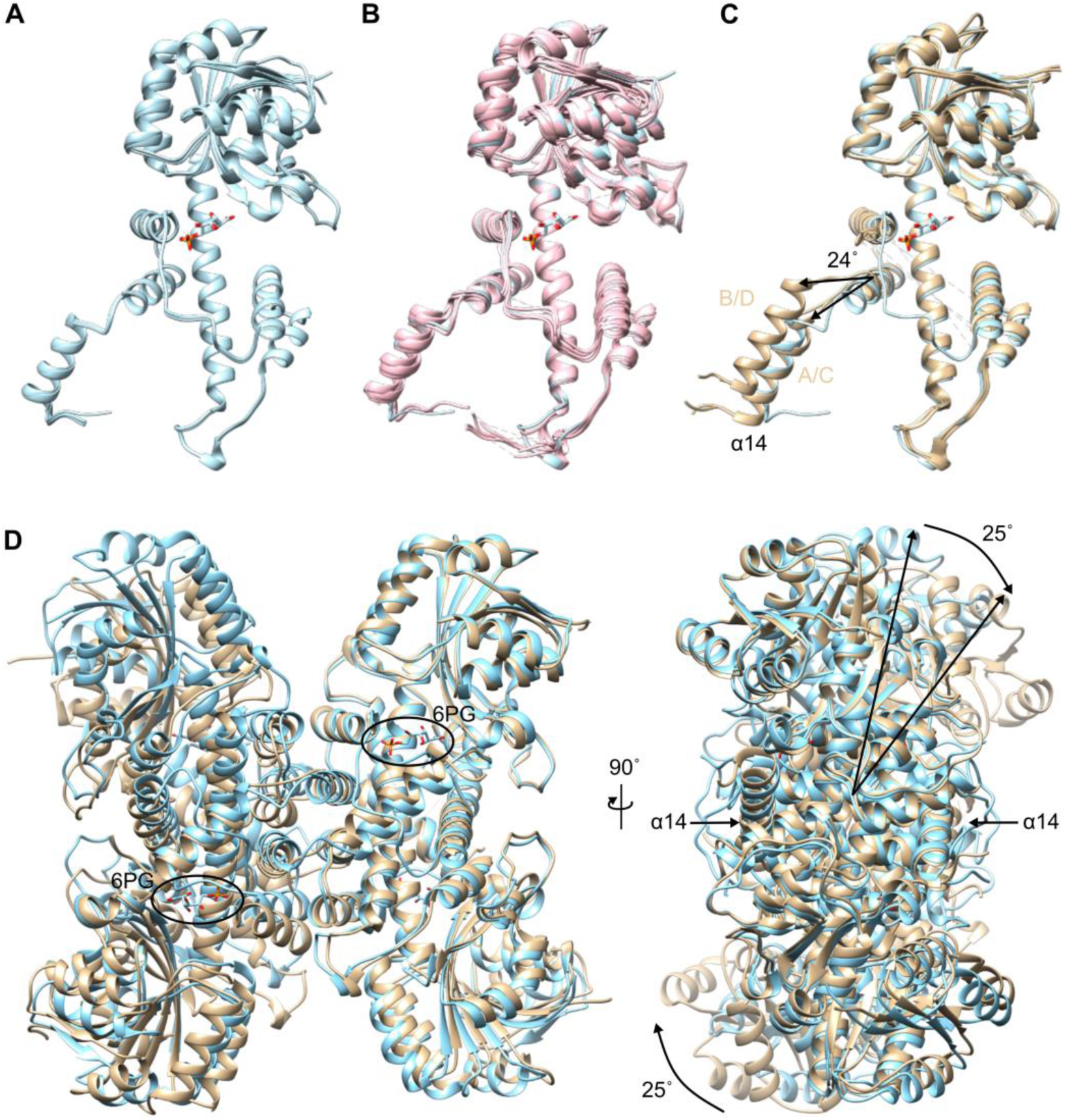
Conformational changes of *Gluconobacter oxydans* 6PGDH in the apo and holo forms. (A) Superposition of the four subunits of holo Go6PGDH (light blue) reveals uniform folding of the N-terminal and C-terminal domains. (B) Structural alignment of the holo Go6PGDH subunit (light blue) with apo subunits from homologous tetrameric 6PGDHs: *Geobacter metallireducens* (*Gm*6PGDH, pink), *Gluconacetobacter diazotrophicus* (*Dz*6PGDH, brown), and Mycobacterium tuberculosis (*Mt*6PGDH, gray). (C) A comparison of the holo (light blue) and apo (light brown) forms of Go6PGDH reveals a 24° outward rotation of the α14 helix in subunits B and D, with subunits A and C remaining aligned. The rotation is centered on the α13 helix. (D) Superimposed apo (brown) and holo (blue) tetrameric structures. In the apo form, a 25° inter-dimeric rotation leads to an expanded quaternary structure. 6PG binding induces tetrameric compaction, reducing inter-dimeric rotation and stabilizing the active site geometry.

**Figure S3.**
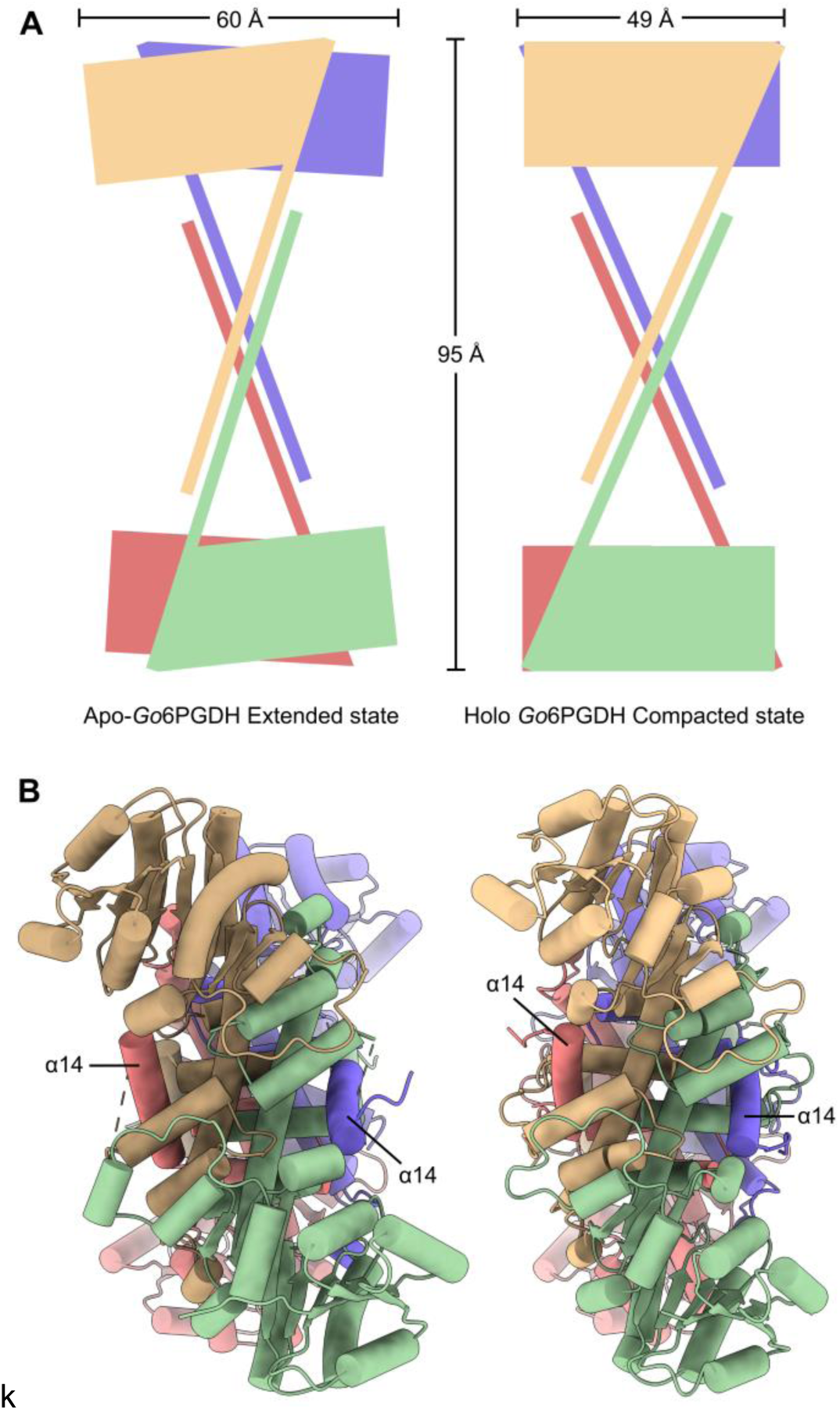
Quaternary compaction of Go6PGDH. (A) Schematic of the apo Go6PGDH tetramer in an extended quaternary state. Colored rectangles depict the relative orientations of the N- and C-terminal domains of each subunit, highlighting increased inter-dimer separation and a wider aperture between opposing C-terminal axes. The maximum inter-dimer distance and lateral spread of the assembly are indicated. (B) Equivalent schematic for the 6-phosphogluconate-bound (holo) *Go*6PGDH tetramer, showing a compact quaternary arrangement.

**Figure S4.**
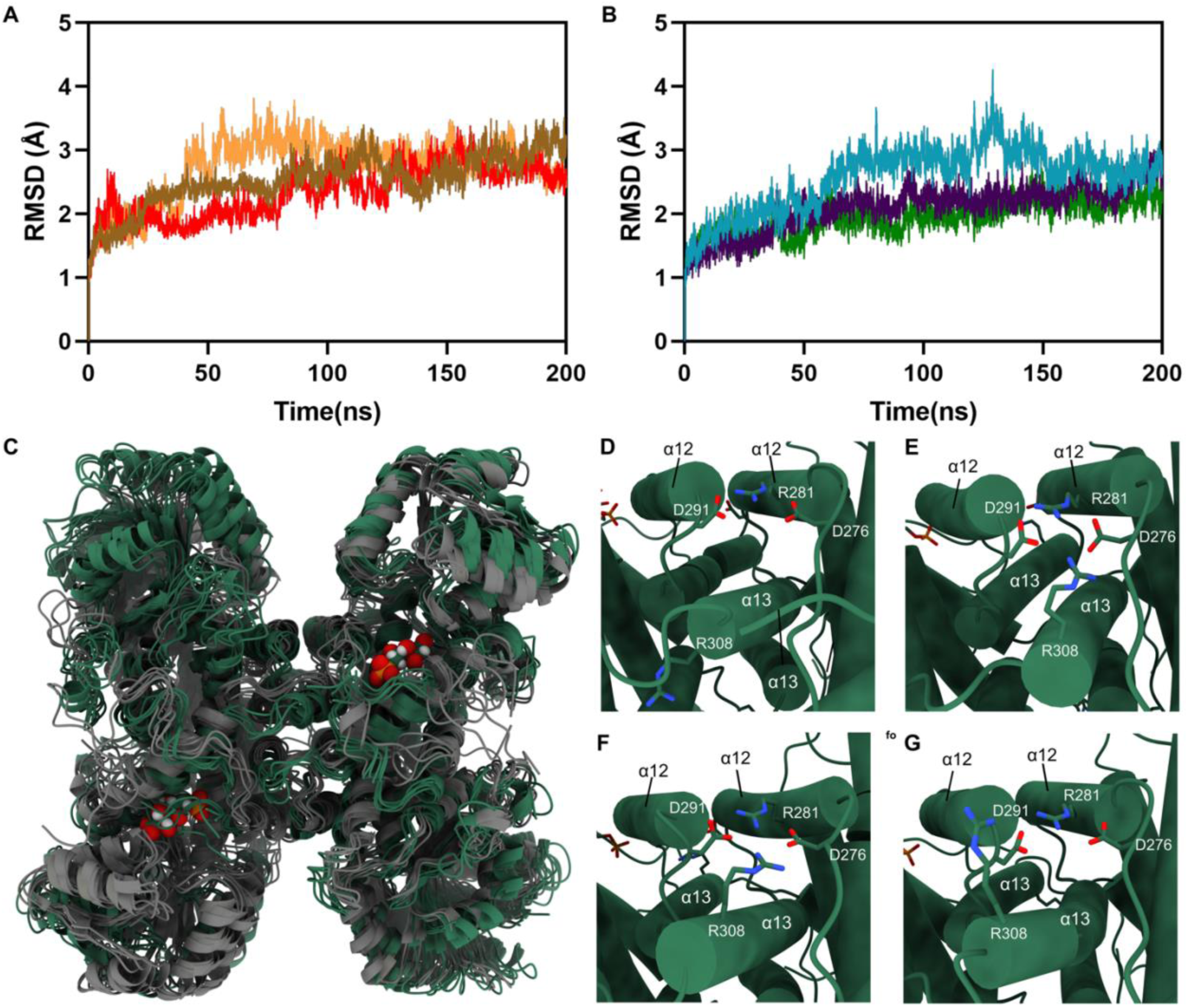
Stability and inter-protomer interface dynamics in apo and 6PG-bound *Go*6PGDH molecular dynamics simulations. (A) Backbone RMSD profiles from three independent 200-ns molecular dynamics simulations of apo *Go*6PGDH. (B) Backbone RMSD profiles from three independent 200-ns simulations of 6PG-bound (holo) *Go*6PGDH. (C) Superposition of representative centroid structures from the five backbone clusters obtained by K-means clustering of the combined apo and holo trajectories. Apo centroids (gray) sample a broader conformational space, whereas holo centroids (green) converge toward a more compact and ordered arrangement around the active site. Bound 6PG molecules are shown as spheres. (D–G) Views of the four inter-protomer interfaces in the apo Go6PGDH tetramer corresponding to centroid 1. Each panel shows the region encompassing helices α12 and α13, highlighting the relative arrangement of residues D276, D291, R281, and R308 that define the inter-protomer contact at each interface. Residues at the interface display variable relative orientations and separations, consistent with a flexible, solvent-exposed interface.

**Supplementary table 1.** Data collection and refinement statistics o 6PG-bound *Go*6PGDH (PDBid: 10GW).

|  |  |
| --- | --- |
| Wavelength | 1.033167 Å |
| Resolution range | 41.61 - 2.0 (2.02 - 2.0) |
| Space group | P 1 |
| Unit cell | 54.118 79.241 86.533 102.17 97 99.4 |
| Total reflections | 318681 (10549) |
| Unique reflections | 90117 (2909) |
| Multiplicity | 3.5 (3.6) |
| Completeness (%) | 97.44 (96.19) |
| Mean I/sigma(I) | 9.56 (1.14) |
| Wilson B-factor | 35.30 |
| R-merge | 0.06872 (1.04) |
| R-meas | 0.08125 (1.223) |
| R-pim | 0.0428 (0.6375) |
| CC1/2 | 0.998 (0.629) |
| CC* | 0.999 (0.879) |
| Reflections used in refinement | 90054 (2905) |
| Reflections used for R-free | 4536 (144) |
| R-work | 0.1935 (0.3682) |
| R-free | 0.2243 (0.4010) |
| Number of non-hydrogen atoms | 10540 |
| macromolecules | 10096 |
| ligands | 68 |
| solvent | 376 |
| Protein residues | 1332 |
| RMS(bonds) | 0.005 |
| RMS(angles) | 0.80 |
| Ramachandran favored (%) | 97.73 |
| Ramachandran allowed (%) | 2.27 |
| Ramachandran outliers (%) | 0.00 |
| Rotamer outliers (%) | 0.39 |
| Clashscore | 5.95 |
| Average B-factor | 69.31 |
| macromolecules | 70.21 |
| ligands | 44.20 |
| solvent | 49.55 |

Statistics for the highest-resolution shell are shown in parentheses.

**Supplementary Table 2.** Clustering analysis of molecular dynamics trajectories of *Go*6PGDH in the apo and 6PG-bound states. Backbone conformations from combined MD trajectories were clustered using the K-means algorithm, yielding five representative clusters for each state. For each cluster, the fraction (%), average distance (Å), and standard deviation are reported.

| Clusters | Apo Go6PGDH |  |  | 6PG-bound Go6PGDH |  |  |
| --- | --- | --- | --- | --- | --- | --- |
|  | % Fraction | Average distance (Å) | Standard deviation (Å) | % Fraction | Average distance (Å) | Standard deviation (Å) |
| Cluster 1 | 33.3 | 2.120 | 0.458 | 33.3 | 1.747 | 0.332 |
| Cluster 2 | 24.7 | 2.056 | 0.436 | 22.6 | 1.696 | 0.286 |
| Cluster 3 | 19.2 | 1.897 | 0.344 | 19.7 | 1.644 | 0.273 |
| Cluster 4 | 14.2 | 1.860 | 0.372 | 13.7 | 1.658 | 0.300 |
| Cluster 5 | 8.6 | 1.893 | 0.372 | 10.7 | 1.614 | 0.275 |

## References

[1] N.J. Kruger, A. Von Schaewen, The oxidative pentose phosphate pathway: structure and organisation, Current Opinion in Plant Biology 6 (2003) 236–246. 10.1016/S1369-5266(03)00039-6.

[2] S. Hanau, J.R. Helliwell, 6-Phosphogluconate dehydrogenase and its crystal structures, Acta Crystallogr F Struct Biol Commun 78 (2022) 96–112. 10.1107/S2053230X22001091.

[3] P. Maturana, E. Tobar-Calfucoy, M. Fuentealba, P. Roversi, R. Garratt, R. Cabrera, Crystal structure of the 6-phosphogluconate dehydrogenase from *Gluconobacter oxydans* reveals tetrameric 6PGDHs as the crucial intermediate in the evolution of structure and cofactor preference in the 6PGDH family, Wellcome Open Res 6 (2021) 48. 10.12688/wellcomeopenres.16572.1.

[4] M.J. Adams, G.H. Ellis, S. Gover, C.E. Naylor, C. Phillips, Crystallographic study of coenzyme, coenzyme analogue and substrate binding in 6-phosphogluconate dehydrogenase: implications for NADP specificity and the enzyme mechanism, Structure 2 (1994) 651–668. 10.1016/S0969-2126(00)00066-6.

[5] Y.-Y. Chen, T.-P. Ko, W.-H. Chen, L.-P. Lo, C.-H. Lin, A.H.-J. Wang, Conformational changes associated with cofactor/substrate binding of 6-phosphogluconate dehydrogenase from *Escherichia coli* and *Klebsiella pneumoniae*: Implications for enzyme mechanism, Journal of Structural Biology 169 (2010) 25–35. 10.1016/j.jsb.2009.08.006.

[6] K. Haeussler, K. Fritz-Wolf, M. Reichmann, S. Rahlfs, K. Becker, Characterization of *Plasmodium falciparum* 6-Phosphogluconate Dehydrogenase as an Antimalarial Drug Target, Journal of Molecular Biology 430 (2018) 4049–4067. 10.1016/j.jmb.2018.07.030.

[7] S. Cameron, V.P. Martini, J. Iulek, W.N. Hunter, Geobacillus stearothermophilus 6-phosphogluconate dehydrogenase complexed with 6-phosphogluconate, Acta Crystallogr F Struct Biol Cryst Commun 65 (2009) 450–454. 10.1107/S1744309109012767.

[8] W. He, Y. Wang, W. Liu, C.-Z. Zhou, Crystal structure of *Saccharomyces cerevisiae* 6-phosphogluconate dehydrogenase Gnd1, BMC Struct Biol 7 (2007) 38. 10.1186/1472-6807-7-38.

[9] Y. Wang, X. Ren, T. Li, D. Su, R. Zhang, Crystal structure and function analysis of 6-phosphogluconate dehydrogenase in *Mycobacterium tuberculosis*, Biochemical and Biophysical Research Communications 731 (2024) 150390. 10.1016/j.bbrc.2024.150390.

[10] W. Kabsch, XDS, Acta Crystallogr D Biol Crystallogr 66 (2010) 125–132. 10.1107/S0907444909047337.

[11] P.R. Evans, G.N. Murshudov, How good are my data and what is the resolution?, Acta Crystallogr D Biol Crystallogr 69 (2013) 1204–1214. 10.1107/S0907444913000061.

[12] L. Potterton, J. Agirre, C. Ballard, K. Cowtan, E. Dodson, P.R. Evans, H.T. Jenkins, R. Keegan, E. Krissinel, K. Stevenson, A. Lebedev, S.J. McNicholas, R.A. Nicholls, M. Noble, N.S. Pannu, C. Roth, G. Sheldrick, P. Skubak, J. Turkenburg, V. Uski, F. Von Delft, D. Waterman, K. Wilson, M. Winn, M. Wojdyr, *CCP* 4 *i* 2: the new graphical user interface to the *CCP* 4 program suite, Acta Crystallogr D Struct Biol 74 (2018) 68–84. 10.1107/S2059798317016035.

[13] A. Vagin, A. Teplyakov, Molecular replacement with *MOLREP*, Acta Crystallogr D Biol Crystallogr 66 (2010) 22–25. 10.1107/S0907444909042589.

[14] P. Emsley, K. Cowtan, *Coot*: model-building tools for molecular graphics, Acta Crystallogr D Biol Crystallogr 60 (2004) 2126–2132. 10.1107/S0907444904019158.

[15] P.D. Adams, P.V. Afonine, G. Bunkóczi, V.B. Chen, I.W. Davis, N. Echols, J.J. Headd, L.-W. Hung, G.J. Kapral, R.W. Grosse-Kunstleve, A.J. McCoy, N.W. Moriarty, R. Oeffner, R.J. Read, D.C. Richardson, J.S. Richardson, T.C. Terwilliger, P.H. Zwart, PHENIX: a comprehensive Python-based system for macromolecular structure solution, Acta Crystallogr D Biol Crystallogr 66 (2010) 213–221. 10.1107/S0907444909052925.

[16] V.B. Chen, W.B. Arendall, J.J. Headd, D.A. Keedy, R.M. Immormino, G.J. Kapral, L.W. Murray, J.S. Richardson, D.C. Richardson, MolProbity: all-atom structure validation for macromolecular crystallography, Acta Crystallogr D Biol Crystallogr 66 (2010) 12–21. 10.1107/S0907444909042073.

[17] M.A. Larkin, G. Blackshields, N.P. Brown, R. Chenna, P.A. McGettigan, H. McWilliam, F. Valentin, I.M. Wallace, A. Wilm, R. Lopez, J.D. Thompson, T.J. Gibson, D.G. Higgins, Clustal W and Clustal X version 2.0, Bioinformatics 23 (2007) 2947–2948. 10.1093/bioinformatics/btm404.

[18] S. Guindon, J.-F. Dufayard, V. Lefort, M. Anisimova, W. Hordijk, O. Gascuel, New Algorithms and Methods to Estimate Maximum-Likelihood Phylogenies: Assessing the Performance of PhyML 3.0, Systematic Biology 59 (2010) 307–321. 10.1093/sysbio/syq010.

[19] T. Müller, M. Vingron, Modeling Amino Acid Replacement, Journal of Computational Biology 7 (2000) 761–776. 10.1089/10665270050514918.

[20] V. Lefort, J.-E. Longueville, O. Gascuel, SMS: Smart Model Selection in PhyML, Molecular Biology and Evolution 34 (2017) 2422–2424. 10.1093/molbev/msx149.

[21] W. Humphrey, A. Dalke, K. Schulten, VMD: Visual molecular dynamics, Journal of Molecular Graphics 14 (1996) 33–38. 10.1016/0263-7855(96)00018-5.

[22] B. Webb, A. Sali, Comparative Protein Structure Modeling Using MODELLER, CP in Bioinformatics 54 (2016). 10.1002/cpbi.3.

[23] M.H.M. Olsson, C.R. Søndergaard, M. Rostkowski, J.H. Jensen, PROPKA3: Consistent Treatment of Internal and Surface Residues in Empirical p *K*_a_ Predictions, J. Chem. Theory Comput. 7 (2011) 525–537. 10.1021/ct100578z.

[24] D.A. Case, H.M. Aktulga, K. Belfon, D.S. Cerutti, G.A. Cisneros, V.W.D. Cruzeiro, N. Forouzesh, T.J. Giese, A.W. Götz, H. Gohlke, S. Izadi, K. Kasavajhala, M.C. Kaymak, E. King, T. Kurtzman, T.-S. Lee, P. Li, J. Liu, T. Luchko, R. Luo, M. Manathunga, M.R. Machado, H.M. Nguyen, K.A. O’Hearn, A.V. Onufriev, F. Pan, S. Pantano, R. Qi, A. Rahnamoun, A. Risheh, S. Schott-Verdugo, A. Shajan, J. Swails, J. Wang, H. Wei, X. Wu, Y. Wu, S. Zhang, S. Zhao, Q. Zhu, T.E. Cheatham, D.R. Roe, A. Roitberg, C. Simmerling, D.M. York, M.C. Nagan, K.M. Merz, AmberTools, J. Chem. Inf. Model. 63 (2023) 6183–6191. 10.1021/acs.jcim.3c01153.

[25] D.A. Case, T.E. Cheatham, T. Darden, H. Gohlke, R. Luo, K.M. Merz, A. Onufriev, C. Simmerling, B. Wang, R.J. Woods, The Amber biomolecular simulation programs, J Comput Chem 26 (2005) 1668–1688. 10.1002/jcc.20290.

[26] C. Tian, K. Kasavajhala, K.A.A. Belfon, L. Raguette, H. Huang, A.N. Migues, J. Bickel, Y. Wang, J. Pincay, Q. Wu, C. Simmerling, ff19SB: Amino-Acid-Specific Protein Backbone Parameters Trained against Quantum Mechanics Energy Surfaces in Solution, J. Chem. Theory Comput. 16 (2020) 528–552. 10.1021/acs.jctc.9b00591.

[27] D.R. Roe, T.E. Cheatham, PTRAJ and CPPTRAJ: Software for Processing and Analysis of Molecular Dynamics Trajectory Data, J. Chem. Theory Comput. 9 (2013) 3084–3095. 10.1021/ct400341p.

[28] P.D. Sarmiento-Pavía, A. Rodríguez-Hernández, A. Rodríguez-Romero, M.E. Sosa-Torres, The structure of a novel membrane-associated 6-phosphogluconate dehydrogenase from *Gluconacetobacter diazotrophicus* ( *Gd* 6PGD) reveals a subfamily of short-chain 6PGDs, The FEBS Journal 288 (2021) 1286–1304. 10.1111/febs.15472.

[29] K. Montin, C. Cervellati, F. Dallocchio, S. Hanau, Thermodynamic characterization of substrate and inhibitor binding to *Trypanosoma brucei* 6-phosphogluconate dehydrogenase, The FEBS Journal 274 (2007) 6426–6435. 10.1111/j.1742-4658.2007.06160.x.

[30] T. Hitosugi, L. Zhou, S. Elf, J. Fan, H.-B. Kang, J.H. Seo, C. Shan, Q. Dai, L. Zhang, J. Xie, T.-L. Gu, P. Jin, M. Alečković, G. LeRoy, Y. Kang, J.A. Sudderth, R.J. DeBerardinis, C.-H. Luan, G.Z. Chen, S. Muller, D.M. Shin, T.K. Owonikoko, S. Lonial, M.L. Arellano, H.J. Khoury, F.R. Khuri, B.H. Lee, K. Ye, T.J. Boggon, S. Kang, C. He, J. Chen, Phosphoglycerate Mutase 1 Coordinates Glycolysis and Biosynthesis to Promote Tumor Growth, Cancer Cell 22 (2012) 585–600. 10.1016/j.ccr.2012.09.020.

[31] R. Liu, W. Li, B. Tao, X. Wang, Z. Yang, Y. Zhang, C. Wang, R. Liu, H. Gao, J. Liang, W. Yang, Tyrosine phosphorylation activates 6-phosphogluconate dehydrogenase and promotes tumor growth and radiation resistance, Nat Commun 10 (2019) 991. 10.1038/s41467-019-08921-8.

[32] I. Berneburg, M. Stumpf, A.-S. Velten, S. Rahlfs, J. Przyborski, K. Becker, K. Fritz-Wolf, Structure of Leishmania donovani 6-Phosphogluconate Dehydrogenase and Inhibition by Phosphine Gold(I) Complexes: A Potential Approach to Leishmaniasis Treatment, IJMS 24 (2023) 8615. 10.3390/ijms24108615.

[33] C. Phillips, J. Dohnalek, S. Gover, M.P. Barrett, M.J. Adams, A 2.8 Å Resolution Structure of 6-Phosphogluconate Dehydrogenase from the Protozoan Parasite *Trypanosoma brucei*: Comparison with the Sheep Enzyme Accounts for Differences in Activity with Coenzyme and Substrate Analogues, Journal of Molecular Biology 282 (1998) 667–681. 10.1006/jmbi.1998.2059.

